# Intravital Deep-Tumor Single-Beam 2-, 3- and 4-Photon Microscopy

**DOI:** 10.1101/2020.09.29.312827

**Authors:** Gert-Jan Bakker, Sarah Weischer, Judith Heidelin, Volker Andresen, Marcus Beutler, Peter Friedl

## Abstract

Three-photon excitation has recently been introduced to perform intravital microscopy in deep, previously inaccessible layers of the brain. The applicability of deep-tissue three-photon excitation in more heterogeneously structured, dense tissue types remains, however, unclear. Here we show that in tumors and bone, high-pulse-energy low-duty-cycle infrared excitation near 1300 and 1700 nm enables two-up to fourfold increased tissue penetration compared to conventional 2-photon excitation. Using a single laser line, simultaneous 2-, 3- and 4-photon processes are effectively induced, enabling the simultaneous detection of blue to far-red fluorescence together with second and third harmonic generation. This enables subcellular resolution at power densities in the focus that are not phototoxic to live cells and without color aberration. Thus, infrared high-pulse-energy low-duty-cycle excitation advances deep intravital microscopy in strongly scattering tissue and, in a single scan, delivers rich multi-parameter datasets from cells and complex organ structures.

## Introduction

Multiphoton microscopy enables studies of the physiology and malfunction of live cells in multicellular organisms^1,2^. Using 2-photon excitation in the near-infrared range, tissue penetration is limited to few tens to hundreds of micrometers, due to light scattering and out-of-focus excitation^3,4^. Recently, this limitation was overcome by 3-photon (3P) microscopy^5,6^, based on low-duty-cycle high-pulse-energy infrared (heIR) excitation^7^. HeIR excitation enables non-invasive detection of brain structures and neuronal calcium signaling beyond 1-mm tissue penetration^8–10^ and direct multimodal visualization of cell morphology and metabolites near the tumor-stroma interface^11,12^. However, the added value of 3P intravital microscopy in dense and heterogeneously organized parenchymatous tissues remains unclear. We here demonstrate that heIR excitation at the spectral windows near 1300 and 1700 nm enables two- to fourfold improved imaging depth in strongly scattering tissues, including tumors and thick bone.

## Results and Discussion

### Setup characterization

Deep 3P microscopy depends on high-energy (within nJ range) excitation with sub-100 fs pulses^5,11^. We applied excitation at 1300 or 1650 nm, to accommodate the spectral range with minimum attenuation of excitation light by combined water absorption and tissue scattering^7^. Excitation was achieved using an optical parametric amplifier (OPA) running at a 1 MHz repetition rate (Supplementary Figure 1a-c and Methods). The pulse lengths under the objective lens were 53 fs for 1300 nm and 89 fs for 1650 nm. Lateral and axial resolutions were 0.721+/-0.014, 2.99+/-0.02 μm for 1300 nm and 0.76+/-0.07, 3.0+/-0.7 μm for 1650 nm (Supplementary Figure 1d). Using 1650 nm excitation to visualize the mouse brain, an imaging depth beyond the cortical region (> 1 mm) and a characteristic attenuation length of 336 μm was obtained (Supplementary Figure 2), similar to independent results^7,10,13^. To ensure compatibility of heIR excitation with longitudinal imaging of in vivo tumor models, saline was used as immersion liquid. Deuterium Oxide, which is being used in brain imaging with impermeable imaging windows annealed to the skull bone^14,15^, is toxic to live tissues^16^, and therefore not compatible with removable intravital microscopy windows inserted in the mouse skin. Compared to Deuterium Oxide, water immersion absorbed approximately 2/3 of the 1650 nm excitation power before the sample surface is reached (Supplementary Figure 3)^17^. Because of the peak power of 87 nJ under the objective, the available excitation energy (Supplementary Figure 1a-b) was sufficient to overcome this additional water absorption, reaching focal planes deep inside the sample.

### Simultaneous 2-, 3- and 4-photon microscopy with a single laser line

When applied to multicolor-fluorescent HT-1080 sarcoma tumors in the deep dermis, excitation at 1300 nm and 1650 nm generated multiple distinct signals, including fluorescent proteins (eGFP, TagRFP, mCherry), a far-red chemical compound for vascular labeling (Dextran70-AlexaFluor680 [AF680]) and blue fluorescence (Hoechst 33342), together with second harmonics generation (SHG) and third harmonics generation (THG) signals (Figure 1a-c, Supplementary Movie 1). To understand the nonlinear processes underlying this broad-range excitation from blue (Hoechst) up to far-red (AF680) fluorophores with a single wavelength, we quantified the dependence between excitation energy and the detected signals and fitted the data to a power law (Figure 1e, Supplementary Figure 4 and Supplementary Table 1). SHG and THG signals, which served as a control, showed respective second- and third-order processes^18^. Excited at 1650 nm, TagRFP, mCherry and AF680 showed a cubic dependence, while Hoechst and eGFP followed a quartic dependence on excitation power below the fluorescence saturation limit, consistent with respective third- and fourth-order processes, as described^18^. This shows that simultaneous 2-, 3- and 4-photon excitations (2PE, 3PE, 4PE) were achieved using 1650 nm excitation, and this resulted in up to 6-channel images in a single scan. The single-pass excitation occurs through a single wavelength and, thus, lacks wavelength dependent aberration.

**Figure 1.**
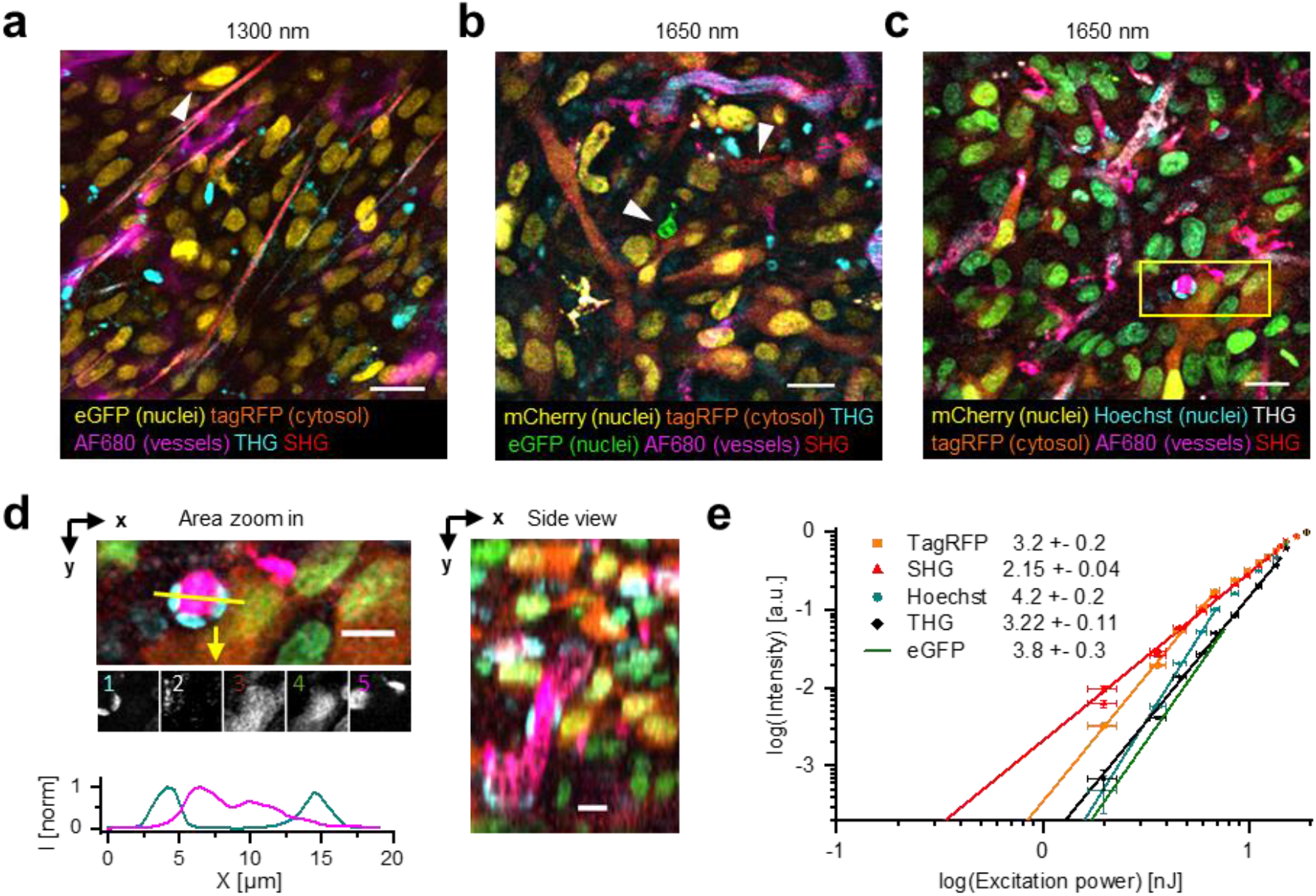
Microscopy with simultaneous 2-, 3- and 4 photon processes excited in fluorescent skin tumor xenografts *in vivo*. Representative images were selected from median-filtered (1 pixel) z-stacks, which were taken in the center of fluorescent tumors through a dermis imaging window. a) Excitation at 1300 nm (OPA) in day-10 tumor at 145 μm imaging depth with a calculated 3.3 nJ pulse energy at the sample surface, 24 μs pixel integration time and 0.36 μm pixel size. For calculation of pulse energy at the sample surface see Figure S3. b) Excitation at 1650 nm (OPA) in day-13 tumor at 30 μm depth with a calculated 6.3 nJ pulse energy at the sample surface, 12 μs pixel integration time and 0.46 μm pixel size. c) Excitation at 1650 nm (OPA) in day-14 tumor at 85 μm depth, with a calculated 5.4 nJ pulse energy at the sample surface, 12 μs pixel integration time and 0.46 μm pixel size. Cell nuclei containing a mixture of mCherry and Hoechst appear as green. d) Zoomed xy-plane (left) and as an orthogonal (xz-) projection (right) (from c, yellow rectangle). Further zoomed detail (middle panel) in single-channel representation (numbers 1-5: Hoechst, THG, TagRFP, mCherry and AF680). Intensity plot (lower panel) of complementary channels along the yellow line, highlighting the positional precision of simultaneously excited fluorophores AF680 and Hoechst. e) Normalized emission intensity (S) as a function of excitation energy (P), for TagRFP, SHG, Hoechst, THG and eGFP recorded with an excitation wavelength of 1650 nm. Data was fitted with *S(P)* = *A·P^n^*, with *A* the proportional factor and *n* the order of the excitation process (indicated as numbers in legend). For curve fitting, excitation intensities at the sample surface below the threshold of physical damage (14 nJ) for SHG and THG and up to their saturation limit (7.6 nJ, eGFP; 6.9 nJ, TagRFP and Hoechst) for fluorophores were used. Images were acquired at the same position as panel (c), except for the fit line of eGFP, which was retrieved from a different dataset (Figure S4). Bars, 25 μm (a-c); 12.5 μm (d).

### Characterization of phototoxicity and bleaching

We next investigated whether the required power densities caused photobleaching and phototoxicity. For heIR excitation, three distinct types of phototoxicity can compromise biological live-cell samples, including: (i) nonlinear processes in the focus where pulsed excitation energy (expressed in nJ) is concentrated and induces toxic reactive oxygen species and photobleaching^19,20^; (ii) transient temperature rise by water absorption in the focus during pulsed excitation (expressed in nJ) causes thermal damage^21^; and (iii) heating over longer spatial and temporal scales, primarily by absorption of out-of-focus photons (expressed in mW), induces thermal damage in and near the scanned volume^20,22^. Using 2.6 nJ (1300 nm) or 8.8 nJ (1650 nm) excitation energy, no notable decrease of 3PE eGFP signal was observed over 75 minutes of three-dimensional (3D) scanning, while mCherry intensity decreased by 10-20 % after 50 minutes (Supplementary Figure 5a). While this level of photobleaching may be incompatible with scanning at high frame rates, as required for Ca^2+^ imaging in the brain^10^, it was within an acceptable range for monitoring the tumor microenvironment, which typically requires low frame rates (15 min up to days), but large-volume scanning^23,24^. As a readout for cell stress caused by nonlinear processes or transient heating in the focal volume, we recorded the intracellular Ca^2+^ influx of tumor cells in vivo (Figure 2a)^25–27^. During continuous OPA exposure for energies at the sample surface below 2.8 nJ (1300 nm) or 8.7 nJ (1650 nm), the Ca^2+^ signal retained background activity, with occasional spontaneous Ca^2+^ fluctuations (Figure 2b, asterisk; Supplementary Movies 2, 3). Higher excitation energies induced Ca^2+^ signaling in cell subsets (Figure 2b and Supplementary Movies 2, 3, arrowheads). These Ca^2+^ responses differed from the background fluctuations by their steep or gradual increase of signal (Figure 2c, d, arrowheads). At prolonged exposure above the observed thresholds, Ca^2+^ signal induction preceded the onset of burning marks (Figure 2b and Supplementary Movies 2, 3, closed arrowheads) or intravascular blood stasis (Supplementary Figure 5b). Furthermore, to avoid thermal damage induced by heating, we applied average power levels under the imaging objective below 100 mW, which in the brain suffices to limit tissue heating below ~1.8 °C^22,28,29^. Thus, we established a limit for power densities to be used for multimodal excitation in tumors to remain below functional phototoxicity levels and showed that higher doses induced different grades of damage^27,30^.

**Figure 2.**
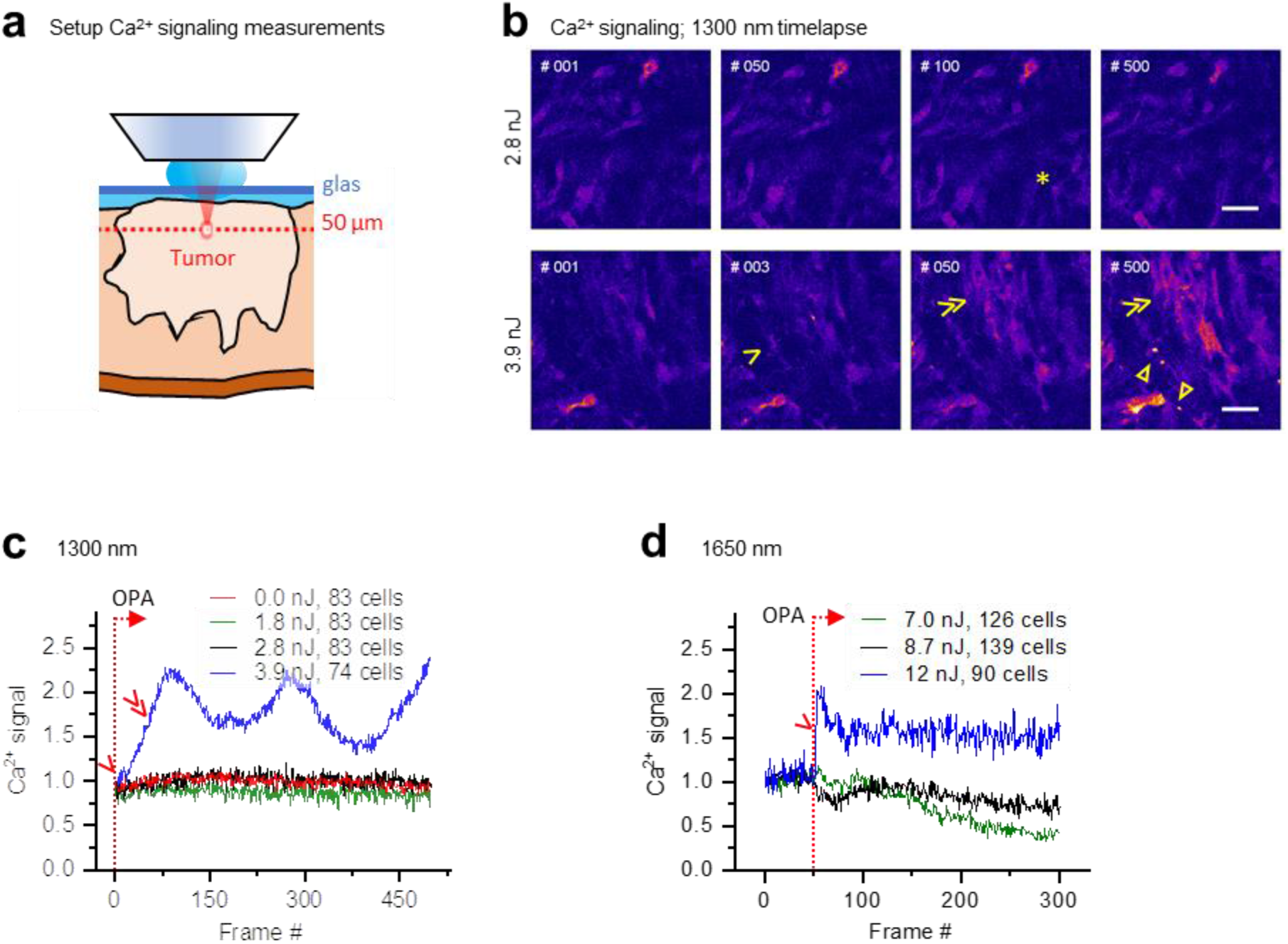
Thresholds of functional phototoxicity at 1300 nm and 1650 nm excitation in tumors *in vivo*. a) Measurement setup. Intradermal B16F10 melanoma expressing GCaMP6 (Ref. ^45^) + H2B-mCherry after 6 (1650 nm) or 11 days (1300 nm) of growth were exposed to OPA laser excitation of increasing dose (50 μm imaging depth). Simultaneously, the Ca^2+^ signal was detected in the same focal plane using low-power Ti:Sa laser excitation at 910 nm (11mW, 0.14 nJ pulse energy). The pixel integration time was 6 μs, pixel size 0.65 or 0.70 μm, and sampling time 1.4 or 1.6 s per frame for excitation at 1300 or 1650 nm, respectively. b) Ca^2+^ signaling in individual cells. Representative frames from the 1300 nm OPA time-series at calculated pulse energies at the sample surface below (2.8 nJ), or above (3.9 nJ) the toxicity threshold. Asterisk, spontaneous, reversible Ca^2+^ signal in single cell, as seen in ~10 of 83 cells in the field of view. Arrowhead, Persistent Ca^2+^ signal starting at frame 2, as seen in 8 of 74 cells. Double arrowheads, multiple cells developing increasing Ca^2+^ signal, present in 45 of 74 cells. Closed arrowheads, burning marks. The frame number is indicated. Bar, 50 μm. c), d) Ca^2+^ signal as a function of time for increasing excitation powers, recorded with an excitation wavelength of 1300 nm (c) or 1650 nm (d). Single and double arrowheads: steep and gradual Ca^2+^ rise, respectively, related to cell populations as described in (b). Emission signal was retrieved by averaging over image area, background subtraction and normalization to the first OPA-excited frame (#1, 1300 nm and #50, 1650 nm, dotted lines). The number of cells per field is indicated.

### Deep-tumor multiparameter microscopy with heIR excitation

We next compared whether heIR excitation at 1300 and 1650 nm provides an advantage for imaging deep tumors regions, with respect to conventional low-pulse-energy high-duty-cycle infrared (lowIR) excitation at 1180 nm using a titanium sapphire / optical parametric oscillator (Ti:Sa/OPO) combination (Figure 3)^26^. To achieve constant emission with increasing tissue penetration, we escalated the excitation power gradually and within the limits of phototoxicity defined above (Figure 3a, grey profiles). The dynamic power range for exciting fluorescence and higher harmonic signals was respective 2.4x or 5.3x higher for 1650 or 1300 nm, compared to 1180 nm. 3PE and 4PE eGFP and TagRFP were detected at depths beyond 400 μm, which improves the imaging depth by ~2-fold compared to 2PE at 1180 nm (Figure 3a) and by 4-fold compared to 2PE in the NIR wavelength range using a Ti:Sa laser^26^. Consistently, multiparameter recordings were achieved inside the tumor at 350 μm depth using excitation at 1650 nm and 1300 nm, but 1180 nm (Figure 3b). In line with an improved depth range, the signal-to-noise ratio (SNR) of 3PE TagRFP outperformed the SNR of 2PE TagRFP at depths beyond 150 μm (Figure 3c). Because H2B-eGFP expression in HT1080 tumors was very high, 3PE eGFP emission reached the highest SNR.

**Figure 3.**
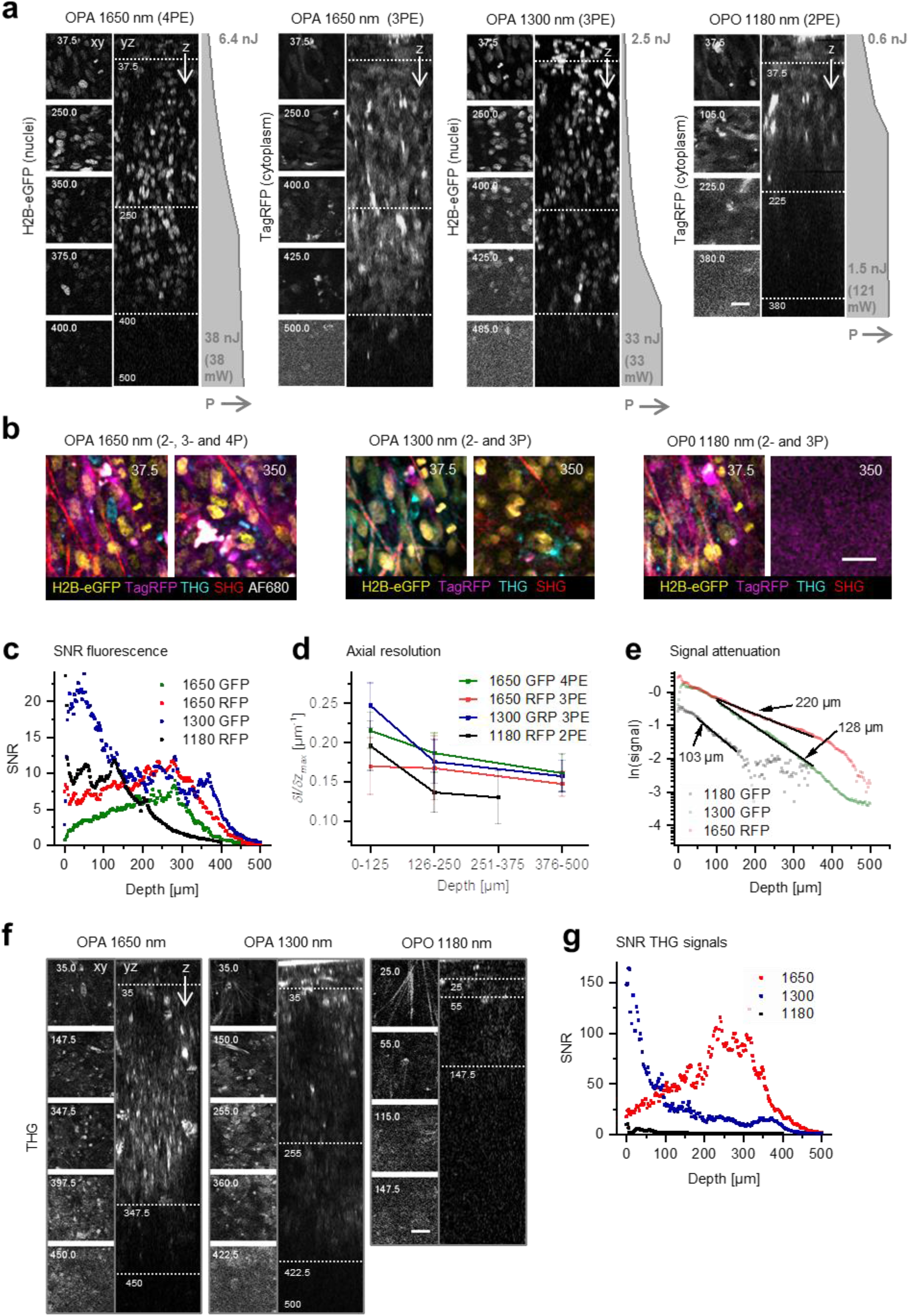
Tissue penetration and resolution of 2-, 3- and 4-photon microscopy in tumors *in vivo*. After 15 days growth, an intradermal HT-1080 sarcoma tumor expressing eGFP and TagRFP was repetitively imaged with OPA and OPO excitation. a) Orthogonal (yz-) views of fluorescent tumor excited at 1650 nm (3PE and 4PE), 1300 nm (3PE) and 1180 nm (2PE). Z-stacks were recorded with increasing power (grey profiles indicating pulse energy at the sample surface as a function of depth). Left, representative (contrast enhanced xy-) images at different depths, represented by the dotted horizontal lines in the yz-views. Specifications: 20 μs pixel integration time, 0.74 μm pixel size, 2.5 μm z-step size. Bar: 25 μm. b) Simultaneous multiparameter microscopy with OPA and OPO excitation, at 37.5 μm and 350 μm depth. Images were processed (median filtered, 1 pixel). Bar: 25 μm. c) SNR of the fluorescent signals as a function of imaging depth, derived from the images shown in (a). d) Axial resolution of the fluorescence signal derived for three depth ranges of the images in panel (a). The steepness of the transition between the normalized intensity of a fluorescent feature and its nonfluorescent surrounding along the z-direction (*(δI/δz)_max_*) was taken as a measure for resolution. For each depth range, the median and standard deviation were calculated over 11-17 fluorescent features per channel. e) Fluorescence signal as a function of imaging depth. The fluorescence intensities derived from (a) were normalized to the square / cubic / quartic of the calculated laser power at the sample surface and to the order of the multiphoton process. For 3PE, characteristic attenuation length *l_e_* was defined as the depth at which the average signal *S* attenuates by *1/e^3^*, where *e* is Euler’s number^7^. *l_e_* (indicated as numbers) were derived from single exponential decay functions (black lines) fitted to the normalized data. f) THG microscopy with OPA and OPO excitation. THG was registered simultaneously and displayed as in (a). Bar: 25 μm. g) SNR of the THG signals in the images shown in panel (e), as a function of imaging depth.

The limits of deep tissue microscopy depend on scattering and aberration of the incident excitation beam^31^. We thus compared how the axial resolution changes with increasing imaging depth and excitation process. For 3PE and 4PE processes, resolution remained high with increasing imaging depth, while the resolution achieved by 2PE declined steeply beyond 125 μm (Figure 3d). Thus, compared to 2PE, 3PE and 4PE improve the resolution in 3D scattering tissue significantly. To address the attenuation of 3PE with increasing tissue penetration in tumors, we measured the fluorescence intensity as a function of increasing scan depth and derived the characteristic attenuation length *l_e_* (Figure 3e). *l_e_* is the mean distance travelled by light before being scattered or absorbed. The decrease of signal remained constant over hundreds of micrometers, indicating that the tumor composition was homogenous over this depth range. When red-shifting the excitation wavelength from 1180 to 1300 or 1650 nm, *l_e_* increased from 103 to 128 or 220 μm, respectively. Similarly, the imaging depth of THG doubled when heIR excitation was used compared to 1180 nm OPO excitation (Figure 3f), in line with a highly increased SNR (Figure 3g).

Compared to lowIR excitation, the gain in resolution and SNR in deep tissue zones with heIR excitation can be attributed to several effects, including: (i) improved localization of the multiphoton effect in the focus^3,5^, (ii) increased *l_e_* at the spectral excitation windows of 1300 and 1700 nm^7,13,32^, and (iii) improved excitation efficiency as a consequence of increased pulse-energy and low laser repetition rate^9^. Through these combined effects, heIR excitation increases the imaging depth by 2-to 4-fold compared to lowIR excitation using OPO- and/or Ti:Sa-based lasers.

### Improved imaging depth of heIR over lowIR in bone

Lastly, we compared lowIR and heIR excitation in tissues of different scatter properties. Bone is strongly light-scattering tissue, yet thin cortical bone such as the mouse skull is amenable to heIR excitation^32,33^. To address whether thick bone can be effectively penetrated by heIR, we performed THG microscopy in an excised ossicle generated in the live mouse^34^. Imaging depth improved by 2-fold (Figure 4a; YZ-projections) and *l_e_* improved by 1.5-fold, comparing 1650 nm heIR versus 1270 nm lowIR excitation (Figure 4b). Subcellular structures were reliably resolved by heIR excitation, including osteocyte lacunae and canaliculi in the cortical bone layer and trabeculae in the bone marrow (Figure 4a; arrowheads). At comparable pulse energies and near the surface (< 105 μm, 1300 versus 1270 nm), the best SNR was obtained with lowIR excitation, taking advantage of its 80-times higher repetition rate and thus increased emission photon flux (Figure 4c, left profiles). However, at greater depth (> 165 μm), the SNR of heIR excitation was superior (Figure 4c, right profiles), consistent with improved maintenance of the excitation power of heIR over lowIR in the focal plane during deep-tissue microscopy. Thus, as in thin bone^33^, heIR excitation improves deep bone microscopy. When comparing the applicability of heIR for tissues with varying attenuation length *l_e_*, including brain, tumor and bone (Figure 4d), the depth gain of THG imaging was approximately 2-fold compared to lowIR excitation and irrespective of tissue type (compare Figure 3f, 4a and Supplementary Figure 2a).

**Figure 4.**
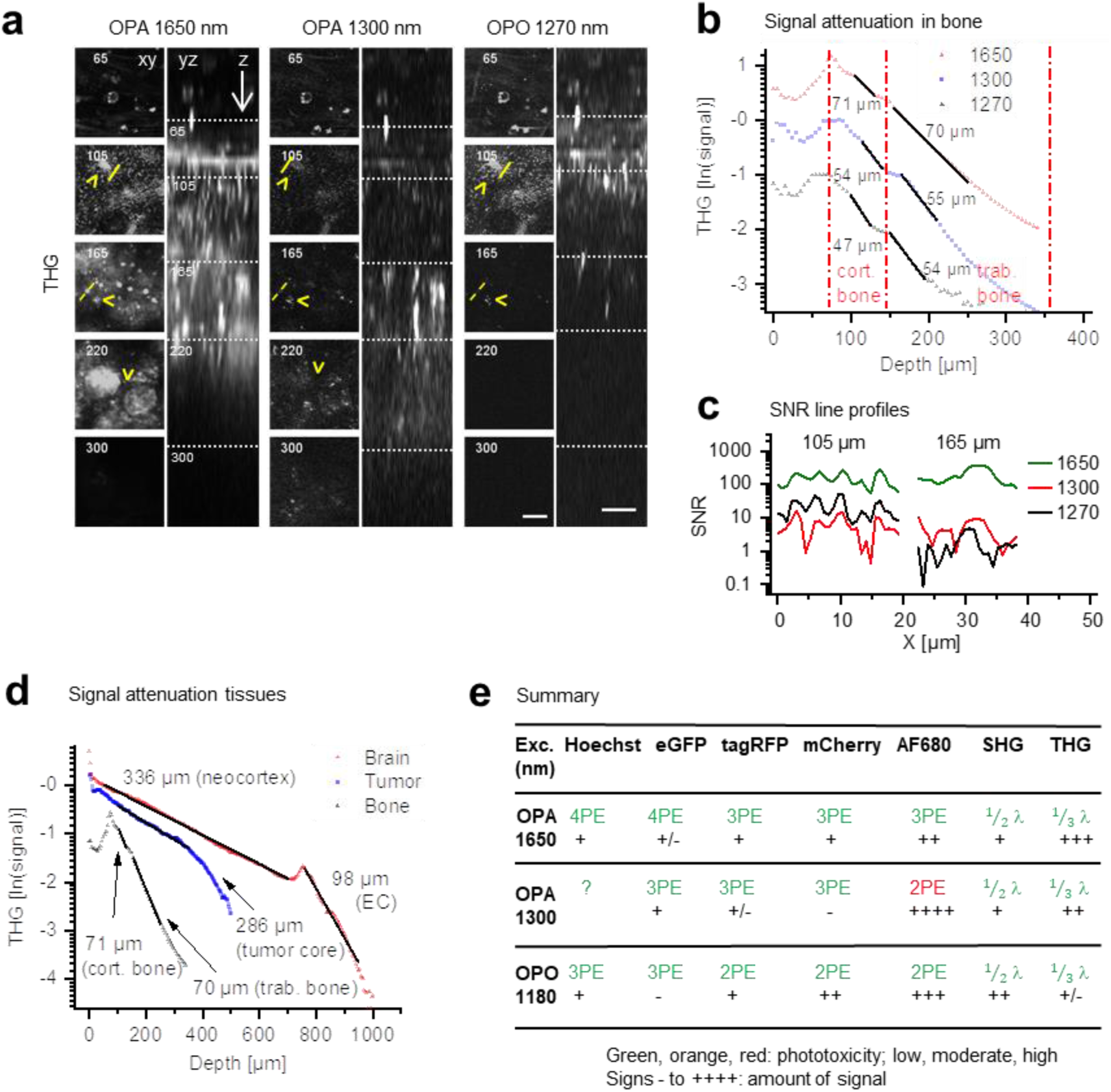
2-, 3- and 4-photon microscopy in scattering tissue samples. a) THG imaging of ex-vivo bone scaffold, with 1650 nm OPA (4.6-32 nJ, left), 1300 nm OPA (1.3-30 nJ, middle) and 1270 nm OPO (1.1-1.9 nJ, right) excitation (pulse energy at the sample surface - increasing with imaging depth to the maximum pulse energy). The xy-images represent the intersections in the yz-images (dotted lines). Specifications: 6 μs pixel integration time, 0.74 μm pixel size, 5 μm z-step size. Bar: 25 μm. b) THG signal as a function of depth. The intensities derived from (a) were normalized (see *methods*) and characteristic attenuation lengths *l_e_* (indicated as numbers) were derived for cortical and trabecular bone layers (black lines). c) SNR derived from THG intensity line profiles drawn over canaliculi ((a), yellow lines z = 105 μm) and other structures (z = 165 μm). d) THG signal in brain (Figure S2), tumor tissue (Figure 3e) and bone (Figure 4a) as a function of imaging depth at 1650 nm excitation. Characteristic attenuation lengths *l_e_* were calculated from discrete tissue layers of varying density/composition (black lines). EC, external capsule. e) Summary of applicability of 2PE, 3PE and 4PE of different fluorophores, based on signal strength and phototoxicity. For each fluorophore the multiphoton process (2-, 3- or 4PE), the amount of signal and the phototoxicity are indicated. For THG and SHG, the emission wavelength and the amount of signal are indicated.

Accumulating evidence suggests that the high pulse energy and average power of heIR excitation is well tolerated by living cells and tissues^32,35^. We calculated the effective excitation pulse energy in the focal plane at the sample surface (Supplementary Figure 3, z = 0) for our experiments, which for 1650 nm was 1.4 to 2.1 times higher, compared to^7,35,36^ and for 1300 nm varied from 0.7 to 1.7 times compared to Refs^8,32^. We showed that phototoxicity and photobleaching were within acceptable range for monitoring dynamic events at time scales typical for the tumor microenvironment (Figure 2 and Supplementary Figure 5)^23,24^. The impact on long-term integrity of cell structure and function require further exploration, including growth, differentiation, and chromatin integrity^20^. Current limitations of heIR excitation include the fluorescence saturation, phototoxicity, and limited emission-photon-rate from the sample^35^, which jointly may compromise recordings with high scan-speed or low fluorescence. Upcoming technical improvements of heIR microscopy include lateral, axial and temporal multiplexing^10^, refined compensation of pulse broadening and detection efficiency^9^ and selective regional scanning^35^, providing means to reduce the photon burden and latent phototoxicity.

## Conclusion

In conclusion, the benefits of heIR excitation observed for brain^7^ also prevail in highly scattering tissues, including thick tumors and bone. Red-shifted excitation by heIR improves both the penetration depth and extends simultaneous multiparameter microscopy of tumors by achieving 4PE fluorescence together with 3PE and multiharmonics^11^. To find the best compromise between the excitation properties of different fluorophores combined with SHG/THG, optimization of settings will require high labeling densities for less efficiently excited fluorophores and empirical choice of wavelength to excite effectively without inducing toxicity (Figure 4e, Supplementary Table 1).

HeIR microscopy provides great potential to advance biomedicine and material sciences. In cancer research, heIR excitation will improve intravital microscopy of understudied regions, including the tumor core and necrosis zones^37^. Beyond cancer, heIR excitation will advance live-tissue microscopy of structurally challenging tissues, including the bone marrow^38^, light-scattering organoids and embryos^20,36^. In addition to fluorescence, the much-improved THG signal, together with SHG, 3PE and 4PE fluorescence, will allow to record cell type and function in a broader morphological context^11,36^, such as biological function of structural interfaces in the tumor microenvironment^39^ and label-free intra-operative histology^40,41^.

## Experimental Section/Methods

### Imaging setup

The setup was based on a customized upright multiphoton microscope (TrimScope II, LaVision BioTec, a Miltenyi Biotec company, Bielefeld, Germany) equipped with two tunable Ti:Sa lasers (Chameleon Ultra I and II, Coherent, California, USA), an OPO (Optical Parametric Oscillator; MPX, APE, Berlin, Germany) and up to 6 PMTs distributed over a 2- and a 4-channel port (Supplementary Figure 1a). The setup was modified to facilitate high-energy, low repetition rate excitation. A high-power fiber laser (Satsuma HP2, 1030 nm, 20W, 1MHz Amplitude Systèmes, Bordeaux, France) was used to pump an OPA (Optical Parametric Amplifier; AVUS SP, APE), which generated 460 mW or 330 mW at 1300 nm or 1650 nm, respectively. A fixed-distance prism compressor (Femtocontrol, APE), glass block (IR-coated 25 mm ZnSe, for 1650 nm) and an autocorrelator with internal and external detector (Carpe, APE) were included in the optical path to control the pulse length under the objective lens. The beam path further included an adjustable 2:1 telescope (f = 80 mm and f = 40 mm apochromat lenses; IR-coated), a motorized half-wave plate and a glan-laser polarizer to control laser beam diameter, power and polarization. The pulse length under the objective lens and its point-spread-function were optimized for the chosen OPA excitation wavelength by adjustment of the excitation path bulk compression, beam pointing and telescope, such that the objective lens back focal plane was 10 % overfilled. A movable mirror was used to guide either the OPO or the OPA beam into the scanhead, where it was spatially overlaid with the Ti:Sa beam. Mirrors, dichroic mirrors and lenses in the scanhead were carefully selected for high reflectance or transmission in the extended excitation wavelength range. Microscopy was performed using a 25x 1.05 NA water immersion objective lens (XLPLN25XWMP2, Olympus, Tokyo, Japan; transmission of 69 % at 1700 nm, data not shown). The following filter / PMT configurations were used: blue-green emission was split off to a 2-channel port with a 560lp dichroic mirror and a 700SP laser blocker filter, while red emission was split off to a 4-channel port with a 900lp dichroic mirror and an 880SP laser blocker filter. Red emission was first split by a 697sp, then further split by a 605lp and a 750sp dichroic mirror, bandpass filtered with 572/28 (TagRFP) or 593/40 (TagRFP and mCherry), 620/60 (mCherry), 710/75 (AlexaFluor680) and 810/90 (AlexaFluor750, SHG) and detected by alkali, GaAsP or GaAs PMT detectors (H6780-20, H7422A-40 or H7422A-50, Hamamatsu, Hamamatsu city, Japan). For 1180 nm, 1270 nm and 1300 nm excitation, blue-green emission was split by a 506lp dichroic mirror, bandpass filtered with 447/60 (THG) and 525/50 (eGFP) and detected by alkali or GaAsP detectors (H6780-01, H6780-20 or H7422A-40, Hamamatsu). For 1650 nm excitation, blue-green emission was split by a 506lp (more THG signal) or 560lp (more eGFP signal) dichroic mirror, bandpass filtered with 447/60 (Hoechst) or 505/40 (eGFP) and 562/40 (THG) and detected by alkali or GaAsP detectors (Hamamatsu, H6780-01, H6780-20 or H7422A-40). Filters were fabricated by Semrock (Newyork, USA) or Chroma Technology GmbH (Olching, Germany). The setup was equipped with a warm plate (DC60 and THO 60-16, Linkam Scientific Instruments Ltd, Tadworth, UK) and a custom-made objective heater (37 °C) for live cell and in vivo experiments, as described^23^.

### Determination of setup resolution

The point-spread-function was obtained using 0.2 μm multicolor beads (FluoresBrite 0.2um, Cat. 24050, Polysciences Inc., Pensylvania, USA). Beads were washed, suspended in agarose (A4718, 1 %w in 1x phosphate buffered saline. Sigma Aldrich, Missouri, USA) and scanned through a coverglass (18×18mm #1, Menzel-Glaeser, Braunschweig, Germany). Z-stacks of 30 μm depth and 0.5 μm step interval were recorded with 1650 nm (5.5 nJ, sample surface), 1300 nm (3.5 nJ, sample surface) and 910 nm (13 mW) excitation with a 0.24 μm pixel size, 1.0 μs pixel dwell time and a 5- (910 nm) or 10-fold (1300 and 1650 nm) line averaging. Red emission was collected using a 650/100 bandpass filter and a GaAsP detector (specified above). The software PSFj^42^ was used for point-spread-function analysis.

### Intravital imaging procedures

Intravital microscopy of intradermal tumors was performed as described^23^. In brief, the animal was anesthetized (1-2 % isoflurane in O_2_ for up to 4.5 h), and vessels visualized using intravenously injected Dextran70-AlexaFlor680 (20-100 μl, 20mg/ml in saline, C29808, Invitrogen, California, USA). At end point sessions, Hoechst 33342 (14533, 1.1 mg in milliQ, Sigma-Aldrich) was injected intravenously to visualize cell nuclei. To define regions of interest, overview images were obtained using an Olympus XL Fluor 4x/340 objective lens and epifluorescence excitation (X-Cite 120 lamp, Excelitas, Massachusetts, USA; Olympus GFP/RFP filter block and a 2/3” cooled CCD camera) (Supplementary Figure 1e). Prior to multiphoton imaging, the maximum average power under the objective was measured (FieldMaxII-TO power meter with PM2 sensor, resolution 1 mW, Coherent) and the excitation energy at the surface of the sample (Supplementary Figure 3) was adjusted below the found functional toxicity threshold (Figure 2, Supplementary Figure 5). To maintain a constant 3-photon emission over imaging depth, the excitation power was increased, with a maximum of 100 mW under the objective to avoid thermal damage^21,22^. For image acquisition, the pixel dwell time was set to 2 or 4 μs to synchronize with the laser repetition rate, line averaging was set between 1 and 6, pixel size was chosen between 0.46-0.82 μm and the step size of z-stacks was 2.5 or 5 μm.

### Brain imaging ex vivo

At the endpoint, a tumor-bearing 9-week-old C57BL/6J mouse was anesthetized, intravenously injected with Dextran70-AF680 and sacrificed. The brain was excised, placed in a phosphate buffered saline filled container and covered with a #1 microscope cover glass (Menzel-Glaser). Z-stack images were acquired in the neocortex above the hippocampus area, with 12 μs pixel dwell time, pixel size 0.50 μm and 4 μm z-step size. Multiple measurements were performed, to optimize either THG and/or AF680 emission for different depth ranges, for 1650 and 1270 nm excitation wavelengths (Supplementary Table 2). Measurements were combined to generate signal attenuation curves and to compose one image stack with maximized penetration depth.

### Image processing and data representation

Unless stated otherwise, image processing was performed with Fiji/ImageJ, version 1.52n^43^. Part of the datasets contained positional jitter, which was removed with the Image Stabilizer plugin^44^. Unless stated otherwise, Origin 2019 (OriginLab Corporation, Massachusetts, USA) was used for numerical and statistical calculations, data fitting and representation.

### Study of the multiphoton processes underlying multimodal excitation

Excitation power under the objective was calibrated for all used attenuator settings. Images were acquired with stepwise decreasing - increasing excitation power. For bleaching correction, reference images were taken after each image. All images were acquired in one imaging plane, with pixel size 0.74 μm and pixel integration time 6.0 μs. For each channel, individual images were merged into two stacks; one for excitation power and one for bleaching correction. To quantify intensities, bright pixels and background pixels were selected by gating with a manually drawn region of interest and/or by multiplication of the image stack with a binary mask. Masks were created by a combination of median filtering, auto-thresholding and binary erode steps. For the selected pixels, area, mean intensity and standard deviation (resp. *I_mean_*, σ_I_ for bright pixels; *B_mean_*, σ_B_ for background) were quantified along the excitation and bleaching correction stacks. Normalized, bleaching corrected, background subtracted mean intensity (*S*) in relation to excitation power (*P*) was derived as follows: *S(P)* = *F_norm_* ·[*I_mean_(P) − B_mean_(P)*]/ [*I_mean_(P_bleach_) B_mean_(P_bleach_)*] where *F_norm_* is a constant for normalization. To estimate the order of the excitation process (*n*), a power function *S(P)* = *A·P^n^* was fitted to the data, with *A* the proportional factor. The Orthogonal Distance Regression iteration algorithm was applied to include both *P* (measurement inaccuracy) and *S* (linear approximation including pixel noise and normalization) errors in the fitting process. Reduced Chi-Square and adjusted R-Square values were below 2 and above 0.995 respectively. Standard errors were given for *A* and *n*.

### Analysis of fluorescence bleaching

The H2B channel (1300 nm, eGFP or 1650 nm, mCherry) of the 3D + time stack was mean (2) filtered and average projected over the z-axis. Bright pixels in cells were selected by auto-thresholding (1300 nm, Huang or 1650 nm, Iso) in combination with manual selection of a region of interest and their average intensity was obtained. The average background was calculated over the manually selected darkest region of the image stack and subtracted from the cell-based fluorescence signal for every time point, to obtain the background subtracted fluorescence signal as a function of time.

### Signal to noise ratio analysis

The SNR as a function of imaging depth was calculated for every position in the depth stack from the average fluorescence intensity (*I_mean_*) over the brightest 1^st^ (THG signal), 10^th^ (nuclei, eGFP) or 40^th^ (cytosol, TagRFP) percentile of pixels in the median filtered (2) image. As background signal, the average (*B_mean_*) and standard deviation (*σ_B_*) were calculated over a ROI in a dark location of the unfiltered stack. Then, the SNR was calculated as *SNR* = (*I_mean_ - B_mean_*)/σ_B_. The SNR along a line profile was obtained from the intensity values along the line (*I_line_*), as *SNR* = (*I_line_ - B_mean_*)/σ_B_.

### Axial resolution analysis of in vivo data stacks

Pixels in fluorescent cell bodies or nuclei in the z-stack were selected with a fixed-size region of interest, their average intensities were calculated over the region of interest and intensity z-profiles were generated. Intensity z-profiles were normalized (0-1) and their maximum derivatives (*(δI/δz)_max_*) were calculated (custom script, Matlab). Median and standard deviation values were derived over sets of (*(δI/δz)_max_*).

### Attenuation length analysis

The fluorescence or THG signal as a function of imaging depth was quantified from each image slice as the average pixel intensity (*I_mean_*), followed by background subtraction. The background (*B_mean_*) was estimated by averaging all the pixel values of the last frame of the image stack. The normalized signal *S* was derived as follows: *S* = *N·[(I_mean_ - B_mean_)/P^n^]^1/n^*, where *N* is a normalization constant, *n* the order of the multiphoton excitation process and *P* the excitation power at the sample surface, which was calculated from the power under the imaging objective and the imaging depth (Supplementary Figure 3). *S* was fitted with a single exponential function to obtain the characteristic attenuation length *l_e_*: *S(z)* = *A · exp(-z/l_e_)*, where *A* is a proportional constant and *z* the imaging depth.

### Cells and cell culture

Murine B16F10 melanoma cells (ATCC, Virginia, USA) were cultured in RPMI (Gibco) supplemented with 10 % FCS (Sigma-Aldrich), 1 % sodium pyruvate (11360, GIBCO, Massachusetts, USA) and 1 % penicillin and streptomycin (PAA, P11/010) at 37 °C in a humidified 5 % CO_2_ atmosphere. Human HT1080 (ACC315) fibrosarcoma cells (DSMZ, Braunschweig, Germany) were cultured in DMEM (Gibco) supplemented with 10 % FCS (Sigma-Aldrich), 1 % sodium pyruvate (11360, Gibco) and 1 % penicillin and streptomycin (PAA, P11/010) at 37 °C in a humidified 5 % CO_2_ atmosphere. Cell line identity was verified by a SNP_ID Assay (Sequenom, MassArray System, Characterized Cell Line Core facility, MD Anderson Cancer Center, Houston, Texas). Cells were routinely tested for mycoplasma using MycoAltert Mycoplasma Detection Kit (Lonza, Basel, Switzerland). HT1080 cells were lentivirally transduced to stably express the fluorescent proteins eGFP or mCherry tagged to histone 2B and cytoplasmic TagRFP. B16F10 cells were lentivirally transduced to stably express the green fluorescent intracellular calcium sensor GCaMP6 (Ref. ^45^) and mCherry tagged to histone 2B.

### 3D spheroid culture

3D spheroid culture was established as described^46^. Shortly, HT1080 fibrosarcoma cells from subconfluent culture were detached with 2mM EDTA (1 mM) and spheroids containing 1000 cells were formed with the hanging drop method. Aggregated spheroids were embedded into a collagen I solution (non-pepsinized rat-tail collagen type I, final concentration 4 mg/ml, REF 354249, Corning, New York, USA) and transferred into a chambered coverglass prior to polymerization at 37 °C. After polymerization, chambers were filled with culture medium (specified above), incubated overnight at 37 °C in a humidified 5 % CO_2_ atmosphere and sealed prior to microscopy.

### Animal procedures

All animal procedures were approved by the ethical committee on animal experimentation (RU-DEC 2014-031) or the Central Authority for Scientific Procedures on Animals (license: 2017-0042). Handlings were performed at the central animal facility (CDL) of the Radboud University, Nijmegen, in accordance with the Dutch Animal experimentation act and the European FELASA protocol. C57Bl/6J WT mice and BALB/c CAnN.Cg-Foxn1nu were purchased from Charles River, Germany. Before the experiment, mice were housed in IVCM cages at standard housing conditions. Food and water were accessible ad libitum. Dorsal skin-fold chambers (DSFC) were transplanted on 8 week to 24 week-old male mice as described^23^. In short, mice were anesthetized using isoflurane anesthesia (2 % in oxygen), the chamber was mounted on the dorsal skinfold of the mice, one side was surgically removed and a cover glass was used to close the imaging window. Mice received an adequate peri-surgical analgesia using carprofen and buprenorphine. To prevent dislocation and inflammation of the DSFC mice were housed with reduced cage enrichment during the experiment. Mice were housed in a temperature-controlled incubator at 28 °C to facilitate tumor growth. One day after surgery, B16F10 melanoma (0.5×10^5^) or HT1080 fibrosarcoma (2×10^6^) were implanted as single cell suspension into the deep dermis of the mouse using a 30G needle (1 or 2 tumors per mouse). To monitor tumor progression, mice were briefly anesthetized using isoflurane and epifluorescence overview images were taken (Supplementary Figure 1e).

## Supporting information

Supplementary file containing Supplementary Figures, Tables and descriptions of Supplementary Movie files.

## Supporting Information

Supplementary information accompanies the manuscript on the Communications Biology website.

## Data Availability

The microscopy image and analysis data that support the findings of this study will be available in/from Figshare.

## Acknowledgements

We acknowledge Eleonora Dondossola for supplying bone samples; Esther Wagena, Bianca Lemmers-Van de Weem and Mike Peters for expert technical support and assistance in animal experiments; and Mirjam Zegers for critical reading of the manuscript. We thank Amplitude Systèmes for providing a Satsuma HP2 demo system and Lucie Desclaux, Yoann Zaouter and Aurelia Durand for hardware support, and we further thank APE GmbH, Berlin, for providing the AVUS SP demo system. Lastly, we gratefully acknowledge Chris Xu, Emmanuel Beaurepaire, Raluca Niesner, Asylkhan Rakhymzhan and Rafael Kurtz for insightful discussions. This work was supported by the European Research Council (617430-DEEPINSIGHT) and the Cancer Genomics Center (CGC.nl) to PF.

## Author contributions

GJB, VA, JH, MB, PF, instrument design and setup and design of experiments. GJB, SW, acquisition of data and analysis. GJB, PF, interpretation of results and writing the manuscript; all authors corrected the manuscript.

