## Supplementary file containing Supplementary Figures, Tables and descriptions of Supplementary Movie files. for "Intravital Deep-Tumor Single-Beam 2-, 3- and 4-Photon Microscopy"

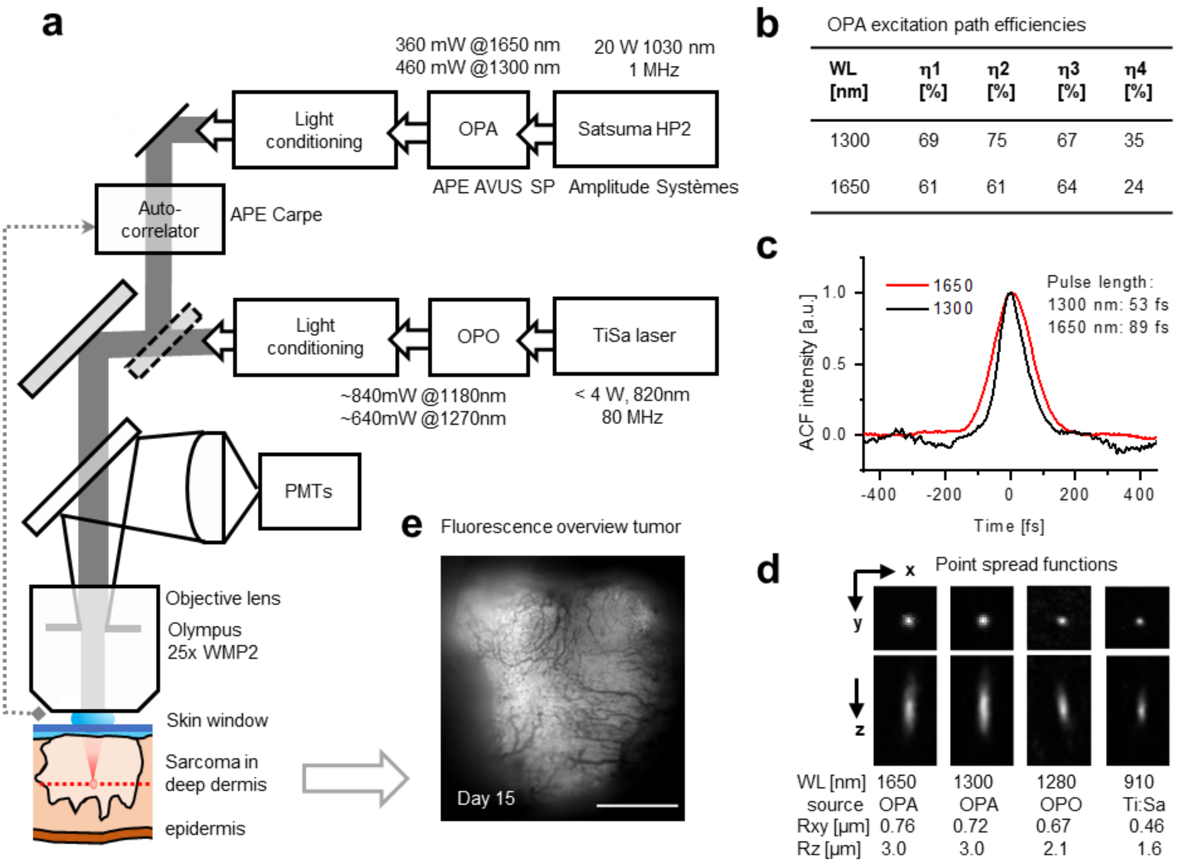


**Supplementary Figure 1.** Microscope setup and excitation parameters. a) Simplified schematics of the setup. Main stages in the excitation path (boxes right to left) including pump laser, OPA or OPO and light conditioning elements. A switchable mirror for selecting OPA or OPO excitation (diagonal dotted box), scanhead (diagonal light grey box), autocorrelator with external detector (dotted line), emission path, detectors (PMTs), objective lens and sample are shown. Light conditioning: pulse compression, attenuation and beam expansion for overfilling of the objective. b) OPA excitation path efficiencies at 1300 and 1650 nm: η1, light conditioning; η2, scanhead; η3, objective lens; η4, complete path. c) Autocorrelation plots of OPA excitation pulses measured under the objective. Pulse lengths were estimated assuming a $Sech^2$ intensity profile. d) Orthogonal and lateral point spread function images of fluorescent beads acquired with 1650 nm OPA, 1300 nm OPA, 1280 nm OPO and 910 nm Ti:Sa excitation. Average lateral (Rxy) and axial (Rz) resolution values and errors were derived over 13, 51, 7 and 67 beads respectively. e) Epi fluorescence overview image of the tumor, made prior to multiphoton measurements used for Figure 3. Fluorescence: TagRFP and eGFP; contrast: blood vessels. Bar: 1 mm.

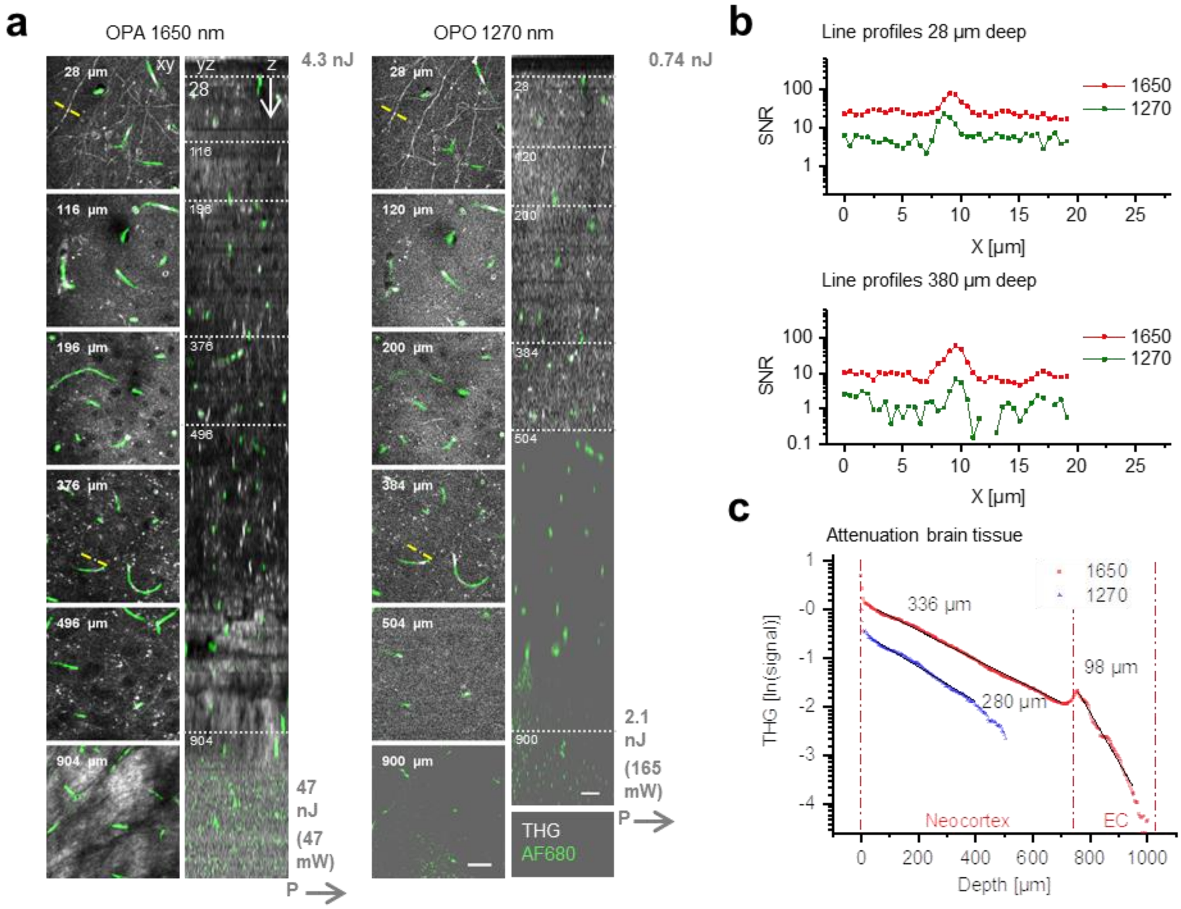


**Supplementary Figure 2.** THG and 3PE microscopy of the mouse neocortex, excited with OPA and OPO excitation, *ex vivo.* a) Orthogonal (yz-) views of THG and fluorescently labeled vessels excited with 1650 nm OPA (left) or 1270 nm OPO (right). Grey profiles indicate pulse energy at the sample surface as a function of imaging depth. Left, representative (contrast enhanced xy-) images at different depths, represented by the dotted horizontal lines in the yz-views. Specifications: 12-71 µs pixel integration time, 0.50 µm pixel size, 4 µm z-step size. Bar: 25 µm. b) SNR derived from THG intensity line profiles drawn over myelinated neuronal features at different depths (panel (a), yellow lines). c) THG signal as a function of depth. The intensities derived from (a) were normalized (see *methods*) and characteristic attenuation lengths $l_e$ (indicated as numbers) were derived for neocortex and EC (external capsule) layers (black lines).

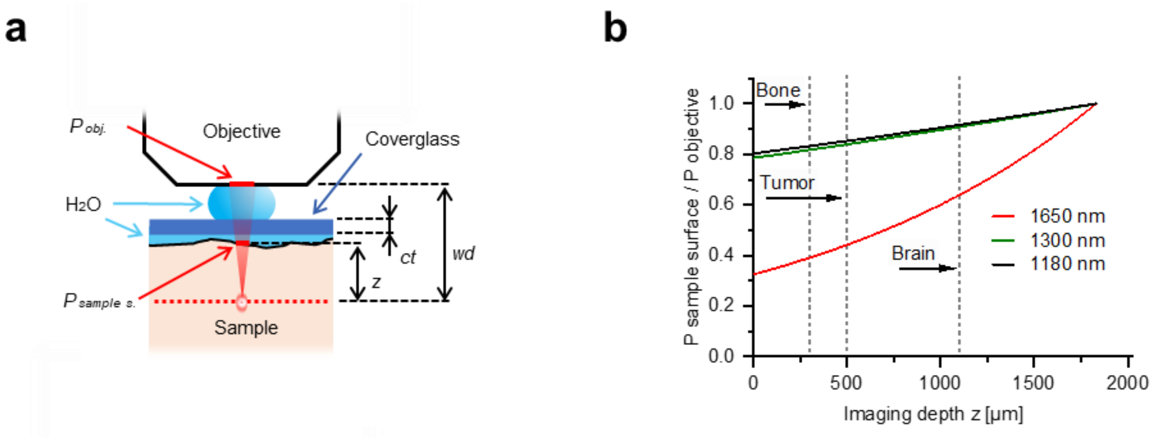


**Supplementary Figure 3.** Attenuation of excitation energy at the sample surface by water immersion. a) The relation between $P_{sample\ surface}/P_{objective}$ and $z$ is given by $P_{sample\ surface}/P_{objective}(z) = e^{\alpha(z-ct+wd)}$, where $\alpha$ is the water absorption at the given wavelength, *ct* the coverglass thickness and *wd* the working distance of the imaging objective. Water absorption $\alpha$ was 6.14, 1.32, 1.2 and 0.07 $cm^{-1}$ at 1650, 1300, 1200 and 900 nm, respectively.[1–3] b) Relative attenuation of excitation light of different wavelengths by water immersion in between the sample and the objective ($P_{sample\ surface}/P_{objective}$) as a function of imaging depth (z). Maximum imaging depth is indicated for bone, tumor and brain tissue (dashed lines).

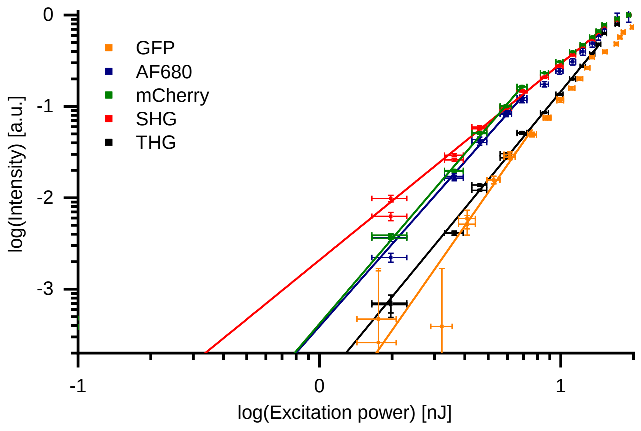


**Supplementary Figure 4.** Two-, 3- and 4-photon processes excited by 1650 and 1300 nm heIR excitation. Normalized emission intensity (*S*) as a function of excitation energy at the sample surface (*P*), for eGFP, AF680, mCherry, SHG and THG, recorded with an excitation wavelength of 1650 nm. Data was fitted with $S(P) = A \cdot P^n$, with *A* the proportional factor and *n* the order of the excitation process.

Maximum excitation energy *P* used for fitting of SHG and THG was below the threshold of physical damage (14 nJ) and up to the saturation limit for fluorophores (7.6 nJ, eGFP; 6.9 nJ, AF680 and mCherry). AF680, mCherry, SHG and THG were acquired during the measurement described in Figure 1e. For eGFP, *S(P)* was derived from 30 images, retrieved from live imaging of human fibrosarcoma spheroids in rattail collagen.

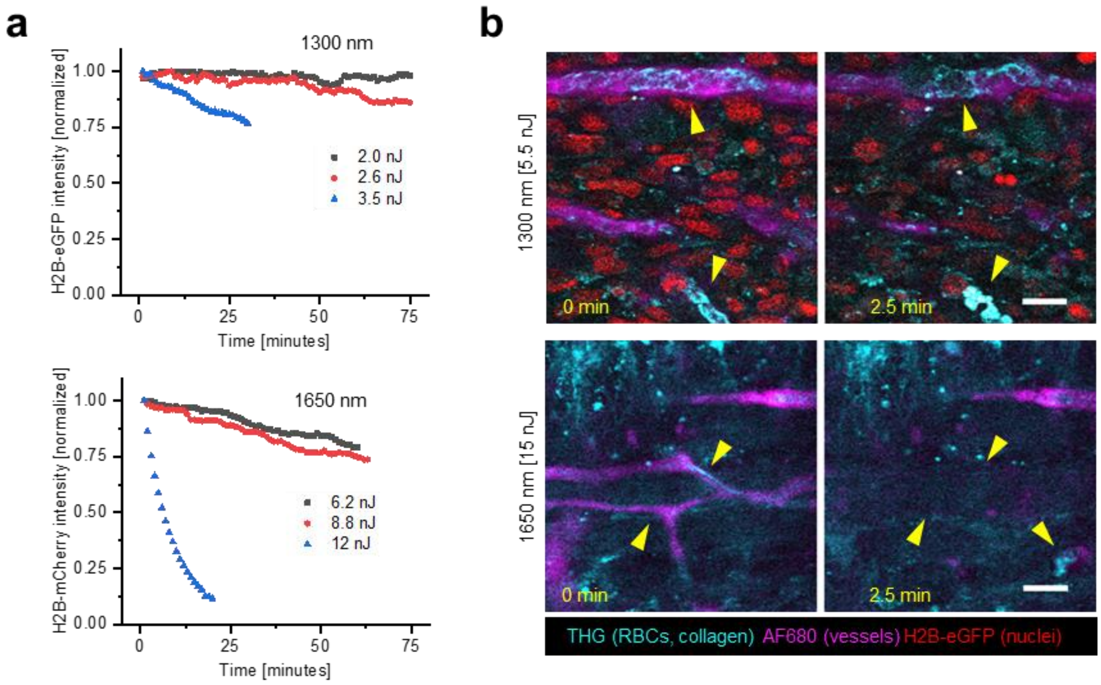


**Supplementary Figure 5.** Scanning-induced fluorophore bleaching and stasis of vascular perfusion. a) Bleaching curves of H2B-eGFP (1300 nm) and H2B-mCherry (1650 nm), retrieved from 3D live cell imaging of human fibrosarcoma spheroids in rattail collagen. Specifications: 0.53 µm pixel size, 5 µm z-step size, 6 µs pixel integration time, 2.3 s per frame and 1 min per stack (11-16 images). Bar: 25 µm. b) Stasis of blood flow in tumor vessels upon 2.5 min of continuous OPA imaging above the functional toxicity threshold. Arrowheads left (0 min): flowing erythrocytes in vessels (THG, stripes and diagonal features), blood vessel labeling (AF680). Arrowheads right (2.5 min): arrested or absent erythrocytes in vessels (THG, round features), blood vessel labeling disappeared. First signs of stasis were observed after 51 or 42 sec (respectively 1300 or 1650 nm, not shown). Specifications: 2 µs pixel integration time, 0.82 µm pixel size, 0.182 s/frame. Fibrosarcoma after 10 days of growth (upper images), melanoma after 6 days of growth (lower images). Bar: 25 µm.

**Supplementary Table 1.** Order of the excitation processes (*n*). Data were derived from datasets described in Figure 1, S4 and from additional independent measurements. Additional measurements, including 1300 nm excitation of eGFP, TagRFP, SHG and THG and 1650 nm excitation of TagRFP and mCherry at 0.5 and 1 MHz repetition rates, were obtained by live imaging of multicellular HT1080 spheroids in rattail collagen. For fitting of data with 1300 nm excitation, the threshold for physical damage was 16 nJ and the saturation limit was 3.1 nJ for eGFP and TagRFP.

| Exc. [nm] | Condition | Hoechst | eGFP | TagRFP | mCherry | AF680 | SHG | THG |
|---|---|---|---|---|---|---|---|---|
| 1650 | *In vivo* | 4.2±0.2 | -- | 3.2±0.2 | 3.1±0.4 | 3.0±0.3 | 2.15±0.04 | 3.2±0.1 |
| 1650 | Spheroids | -- | 3.8±0.3 | 3.0±0.2 | 3.0±0.1 | -- | -- | 3.0±0.2 |
| 1650 | Spheroids[a] | -- | -- | 3.1±0.1 | 3.0±0.2 | -- | -- | -- |
| 1300 | Spheroids | -- | 2.6±0.4 | 2.5±0.4 | | -- | 1.85±0.08 | 2.9±0.3 |

[a] Measured with the excitation source set to a repetition rate of 0.5 MHz.

**Supplementary Table 1.** Experimental parameters for brain measurements optimized for THG and/or AF680 emission. Data were obtained for 1650 heIR or 1270 nm lowIR excitation and different depth ranges. Line averaging (Line av.), 2-or 4-channel emission port (Port), dichroic mirror to split off emission (DM), emission bandpass filter (BP), detector type (PMT) and immersion liquid (Imm.). Rows in the table with the same sequence number (Seq.) were acquired simultaneously.

| Channel / depth / Wavelength [nm] | Line av. | Port | DM | BP | PMT | Imm. | Seq. |
|---|---|---|---|---|---|---|---|
| THG / < 830 µm / 1650 nm | 2 | 2 ch | 525/50 | no | GaAsP | $H_2O$ | 1 |
| THG / > 830 µm / 1650 nm[a] | 6 | 2 ch | 525/50 | no | GaAsP | $H_2O$ | 2 |
| THG / > 830 µm / 1650 nm | 6 | 2 ch | 880lp | no | GaAsP | $D_2O$ | 3 |

| | | | | | | | |
|---|---|---|---|---|---|---|---|
| AF680 / < 830 µm / 1650 nm | 2 | 4 ch | 900lp | 710/75 | GaAs | $H_2O$ | 1 |
| AF680 / > 830 µm / 1650 nm | 6 | 2 ch | 880lp | 710/75 | GaAs | $H_2O$ | 3 |
| THG / < 500 µm / 1270 nm | 3 | 2 ch | 485lp | 417/60 | GaAsP | $H_2O$ | 4 |
| AF680 / < 500 µm / 1270 nm | 3 | 4 ch | 900lp | 710/75 | Alkali | $H_2O$ | 4 |
| AF680 / 500-669 µm / 1270 nm | 1 | 4 ch | 900lp | 710/75 | Alkali | $H_2O$ | 5 |
| AF680 / > 669 µm / 1270 nm | 3 | 4 ch | 900lp | 710/75 | Alkali | $H_2O$ | 6 |

[a)] Solely used for generation of the 1650 nm signal attenuation curve.

**Supplementary Movie 1.** Single-beam excitation at 1650 nm for 2-, 3- and 4 photon microscopy of fluorescent tumor xenograft *in vivo*. Rotation of 3D image stack corresponding to Figure 1c. Stack size: 198 x 198 x 133 $\mu m^3$.

**Supplementary Movie 2.** Functional phototoxicity in B16F10 melanoma tumor expressing the intracellular $Ca^{2+}$ sensor GCaMP6 (Ref. [4]) at 1300 nm OPA laser excitation. The time-course of 500 frames and dose increase corresponds to the Figure 2b, c. Recordings were stopped upon burning marks or whole-field $Ca^{2+}$ signal, indicating general toxicity (5.1 and 5.9 nJ OPA excitation). Arrowheads, onset of $Ca^{2+}$ signaling; closed arrowheads, burning marks. Bar: 50 µm.

**Supplementary Movie 3.** Functional phototoxicity in B16F10 melanoma tumor expressing the intracellular $Ca^{2+}$ sensor GCaMP6 (Ref. [4]) at 1650 nm OPA laser excitation. The time-course of 300 frames and dose increase corresponds to the Figure 2d. Arrowheads, onset of $Ca^{2+}$ signaling; closed arrowhead, burning marks. Bar: 50 µm.
